# The grayling genome reveals selection on gene expression regulation after whole genome duplication

**DOI:** 10.1101/153270

**Authors:** Srinidhi Varadharajan, Simen R. Sandve, Gareth B. Gillard, Ole K. Tørresen, Teshome D. Mulugeta, Torgeir R. Hvidsten, Sigbjørn Lien, Leif Asbjørn Vøllestad, Sissel Jentoft, Alexander J. Nederbragt, Kjetill S. Jakobsen

## Abstract

Whole genome duplication (WGD) has been a major evolutionary driver of increased genomic complexity in vertebrates. One such event occurred in the salmonid family ~80 million years ago (Ss4R) giving rise to a plethora of structural and regulatory duplicate-driven divergence, making salmonids an exemplary system to investigate the evolutionary consequences of WGD. Here, we present a draft genome assembly of European grayling *(Thymallus thymallus)* and use this in a comparative framework to study evolution of gene regulation following WGD. Among the Ss4R duplicates identified in European grayling and Atlantic salmon *(Salmo salar)*, one third reflect non-neutral tissue expression evolution, with strong purifying selection, maintained over ~50 million years. Of these, the majority reflect conserved tissue regulation under strong selective constraints related to brain and neural-related functions, as well as higher-order protein-protein interactions. A small subset of the duplicates has evolved tissue regulatory expression divergence in a common ancestor, which have been subsequently conserved in both lineages, suggestive of adaptive divergence following WGD. These candidates for adaptive tissue expression divergence have elevated rates of protein coding- and promoter-sequence evolution and are enriched for immune- and lipid metabolism ontology terms. Lastly, lineage-specific duplicate divergence points towards underlying differences in adaptive pressures on expression regulation in the non-anadromous grayling versus the anadromous Atlantic salmon.

Our findings enhance our understanding of the role of WGD in genome evolution and highlights cases of regulatory divergence of Ss4R duplicates, possibly related to a niche shift in early salmonid evolution.

## Introduction

Whole genome duplication (WGD) through spontaneous doubling of all chromosomes (autopolyploidization) has played a vital role in the evolution of vertebrate genome complexity (Van de Peer et al. 2009). However, the role of selection in shaping novel adaptations from the redundancy that arises from WGD is not well understood. The idea that functional redundancy arising from gene duplication sparks evolution of novel traits and adaptations was pioneered by Susumu Ohno (Ohno 1970). Duplicate genes that escape loss or pseudogenization are known to acquire novel regulation and expression divergence (Lynch & Conery 2000; Zhang 2003; Conant & Wolfe 2008). Functional genomic studies over the past decade have demonstrated that large-scale duplications lead to the rewiring of regulatory networks through divergence of spatial and temporal expression patterns (Osborn et al. 2003). As changes in gene regulation are known to be important in the evolution of phenotypic diversity and complex trait variation (Carroll 2000; Wray 2003), these post-WGD shifts in expression regulation may provide a substrate for adaptive evolution. Several studies have investigated the genome-wide consequences of WGD on gene expression evolution in vertebrates (e.g. (Sémon & Wolfe 2008; Kassahn et al. 2009; Berthelot et al. 2014; Li et al. 2015; Acharya & Ghosh 2016; Lien et al. 2016; Robertson et al. 2017) and have revealed that a large proportion of gene duplicates have evolved substantial regulatory divergence of which, in most cases, one copy retains ancestral-like regulation (consistent with Ohno’s model of regulatory neofunctionalization). However, to what extent this divergence in expression is linked to adaptation remains to be understood. A major factor contributing to this knowledge gap is the lack of studies that integrate expression data from multiple species sharing the same WGD (Hermansen et al. 2016). Such studies would allow us to distinguish neutral evolutionary divergence in regulation from regulatory changes representing adaptive divergence and those maintained by purifying selection.

Salmonids have emerged as a model for studying consequences of autopolyploidization in vertebrates, owing to their relatively young WGD event (Ss4R, <100MYA) (Ohno 1970; Alexandrou et al. 2013) and ongoing rediploidization (Macqueen & Johnston 2014; Limborg et al. 2016; Lien et al. 2016; Robertson et al. 2017). Directly following autopolyploidization, duplicated chromosomes pair randomly with any of their homologous counterparts resulting in an increased risk of formation of multivalents and consequently production of non-viable aneuploid gametes. Restoring bivalent chromosome pairing is therefore a critical step towards a functional genome post-WGD (Wolfe 2001). This can be achieved through e.g. structural rearrangements that suppress recombination, block multivalent formation, and drive the process of returning to a functional diploid state (i.e. rediploidization). Since the mutational process is stochastic, rediploidization occurs independently for different chromosomes. As a result, the divergence of gene duplicates resulting from WGD (referred to as ohnologs) is also achieved independently for different chromosomes and hence occurs at different rates in various genomic regions. Recent studies on genome evolution subsequent to Ss4R have shown that the rediploidization process temporally overlaps with the species radiation, resulting in lineage-specific ohnolog resolution (LORe) that may fuel differentiation of genome structure and function (Macqueen & Johnston 2014; Robertson et al. 2017). In fact, due to the delayed rediploidization, only 75% of the duplicated genome diverged before the basal split in the Salmonid family ~60 MYA (henceforth referred to as ancestral ohnolog resolution, AORe). Consequently, ~25% of the Ss4R duplicates have experienced independent rediploidization histories after the basal salmonid divergence resulting in the Salmoninae and Thymallinae clades. Interestingly, the species within these two clades have also evolved widely different genome structures, ecology, physiology and life history adaptations (Hendry & Stearns 2004). In contrast to the Thymallus lineage, the species in the subfamily Salmoninae have fewer and highly derived chromosomes resulting from large-scale chromosomal translocations and fusions (Supplementary Figure S1), display extreme phenotypic plasticity, and have evolved the capability of migrating between fresh and saltwater habitats (referred to as anadromy) (Nygren et al. 1971; Phillips & Ráb 2001; Hartley 1987; Ocalewicz et al. 2013; Alexandrou et al. 2013). This unique combination of both shared and lineage-specific rediploidization histories, and striking differences in genome structure and adaptations, provides an ideal study system for addressing key questions about the evolutionary consequences of WGD.

To gain deeper insights into how selection has shaped the evolution of gene duplicates post WGD, we have sequenced, assembled and annotated the genome of the European grayling *(Thymallus thymallus*), a species representative of an early diverging non-anadromous salmonid lineage, Thymallinae. We use this novel genomic resource in a comparative phylogenomic framework with the genome of Atlantic salmon *(Salmo salar*) of the Salmoninae lineage, to address the consequences of Ss4R WGD on lineage-specific rediploidization and selection on ohnolog gene expression regulation.

Our results reveal signatures of adaptive regulatory divergence of ohnologs, strong selective constraints on expression evolution in brain and neural-related genes, and lineage-specific ohnolog divergence. Moreover, diverse biological processes are correlated to differences in evolutionary constraints during the 88–100MY of evolution post-WGD, pointing towards underlying differences in adaptive pressures in non-anadromous grayling and anadromous Atlantic salmon.

## Results

### Genome assembly and annotation

We sequenced the genome of a wild-caught male grayling individual, sampled from the Norwegian river Glomma, using the Illumina HiSeq 2000 platform (Supplementary Table S1 and S2). *De novo* assembly was performed using ALLPATHS-LG (Gnerre et al. 2011), followed by assembly correction using Pilon (Walker et al. 2014), resulting in 24,343 scaffolds with an N50 of 284 Kbp and a total size of 1.468 Gbp (Table 1). The scaffolds represent approximately 85% of the k-mer-based genome size estimate of ~1.8 Gbp. The C-values estimated previously for European grayling are 2.1pg (http://www.genomesize.com/) and 1.9pg (Hartley 1987). To annotate gene structures, we used RNA-Seq data from nine tissues extracted from the sequenced individual. We constructed transcriptome assemblies using both *de novo* and reference-based methods. Repeat masking with a repeat library constructed using a combination of homology and *de* novo-based methods identified and masked approximately 600Mb (~40%) of the assembly and was dominated by class1 DNA transposable elements (Supplementary Table S3 and a repeat landscape in Supplementary Figure S2). Finally, the transcriptome assemblies, the *de novo* identified repeats along with the UniProt proteins (UniProt Consortium 2015) and Atlantic salmon coding sequences (Lien et al. 2016) were utilized in the MAKER annotation pipeline, predicting a total of 117,944 gene models, of which 48,753 protein coding genes were retained based on AED score (Annotation Edit Distance), homology with UniProt and Atlantic salmon proteins or presence of known domains. Assembly completeness was assessed at the gene level by looking for conserved genes using CEGMA and BUSCO. The assembly contains 236 (95.16%) out of 248 conserved eukaryotic genes (CEGs) with 200 (80.65%) complete CEGs. Of the 4,584 BUSCO (database: Actinopterygii, odb9), 4102 complete (89.5%) and 179 (3.9%%) fragmented genes were found in the assembly (Table 1).

**Table 1:** Grayling genome assembly statistics

| Assembly statistics |  | Assembly Validation |  |
| --- | --- | --- | --- |
| **Total size of scaffolds (bp)** | 1,468,519,221 | **Complete CEGMA<sup>a</sup> genes** | 80.65% (200/248) |
| **Number of scaffolds** | 24,369 | **Partial CEGMA genes** | 95.16% (236/248) |
| **Scaffold N50 (bp)** | 283,328 | **Complete BUSCOs<sup>b</sup>** | 4102 (89.5%) |
| **Longest scaffold (bp)** | 2,502,076 | **Complete Duplicated BUSCOs** | 1724 (37.6%) |
| **Total size of contigs (bp)** | 1,278,330,545 | **Fragmented BUSCOS** | 179(3.9%) |
| **Number of contigs** | 216,549 | **Missing BUSCOS** | 303(6.6%) |
| **Contig N50 (bp)** | 11,206 | **Total BUSCOS searched** | 4,584 |

<sup>a</sup> Based on 248 highly Conserved Eukaryotic Genes (CEGS),
<sup>b</sup> Based on a set of 4,584 actinopterygii odb9 Benchmarking Universal Single-Copy Orthologs (BUSCOs)

### Divergent rediploidization rates among the salmonid lineages

Previous studies have suggested that up to 25% of the genome of the most recent common salmonid ancestor was still tetraploid when the grayling and Atlantic salmon lineages diverged (Lien et al. 2016; Robertson et al. 2017). To test this hypothesis, we used a phylogenomic approach to characterize rediploidization following Ss4R in grayling. We inferred 23,782 groups of orthologous genes (i.e. ortholog groups or orthogroups) using gene models from *Homo sapiens* (human)*, Mus musculus* (mouse)*, Danio rerio* (zebrafish)*, Gasterosteus aculeatus* (stickleback), *Oryzias latipes* (medaka), *Esox lucius* (northern pike), *Salmo salar* (Atlantic salmon), *Oncorhynchus mykiss* (rainbow trout) *and Oncorhynchus kisutch* (coho salmon) (Figure 1). These orthogroups were used to infer gene trees. In total, 20,342 gene trees contained WGD events older than Ss4R (Ts3R or 2R) and were further subdivided into smaller sub-groups (i.e. clans, see Methods for details and Supplementary Figure S3). To identify orthogroups with retained Ss4R duplicates, we relied on the high-quality reference genome of Atlantic salmon (Lien et al. 2016). A synteny-aware blast approach (Lien et al. 2016) was first used to identify Ss4R duplicate/ohnolog pairs in the Atlantic salmon genome and this information was used to identify a total of 8,527 gene trees containing high confidence ohnologs originating from Ss4R. Finally, gene trees were classified based on the tree topology into duplicates conforming to LORe and those with ancestrally diverged duplicates following the topology expected under AORe (Figure 2a). In total, 3,367 gene trees correspond to LORe regions (2,403 with a single copy in grayling) and 5,160 correspond to an AORe-like topology. These data were cross-checked with the LORe coordinates suggested by Robertson et al. (Robertson et al. 2017) and genes with LORe-type topologies from non-LORe regions of the genome were discarded. The final set (henceforth referred to as ohnolog-tetrads) consisted of 5,475 gene trees containing Ss4R duplicates from both species (4,735 AORe, 740 LORe). In addition, 482 ortholog sets contained Ss4R duplicates in Atlantic salmon but not in grayling.

**Fig. 1.**
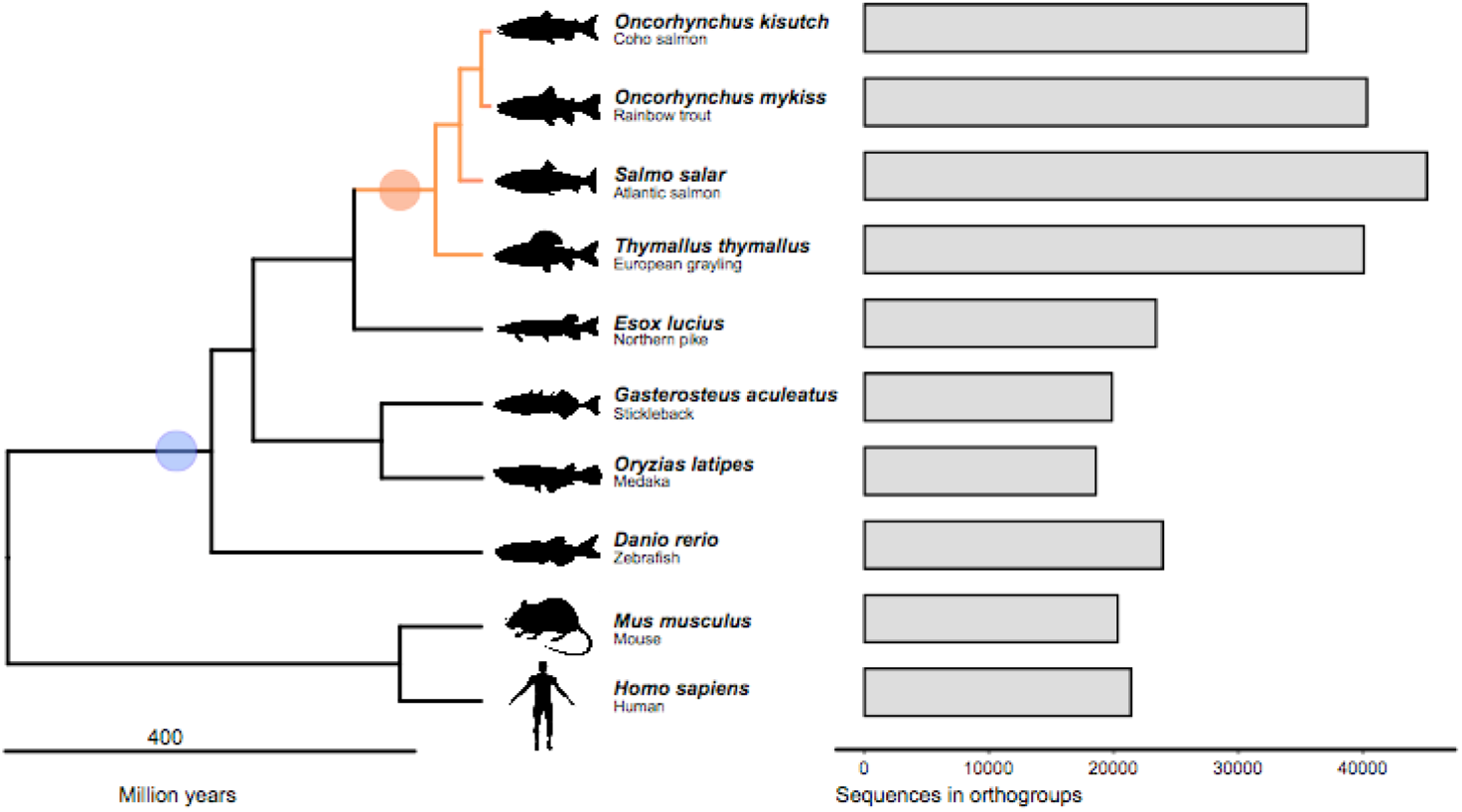
Species and genes in ortholog groups. Left: phylogenetic relationship of species used for constructing ortholog groups and gene trees. The blue circle indicates the 3R-WGD event while the Ss4R event is indicated with an orange circle. Right: number of genes

To identify regions of ancestral and lineage-specific rediploidization in the grayling genome, we assigned genes from gene trees that contained Ss4R duplicates to genomic positions on the Atlantic salmon chromosomes (Figure 2b). In Atlantic salmon, several homeologous chromosome arms (2p-5q, 2q-12qa, 3q-6p, 4p-8q, 7q-17qb, 11qa-26, 16qb-17qa) have previously been described as homeologous regions under delayed rediploidization (Lien et al. 2016; Robertson et al. 2017) (indicated in Figure 2b as red and blue ribbons). Interestingly, for the homeologous LORe regions 2q-12qa, 4p-8q and 16qb-17qa in Atlantic salmon, we identified only one orthologous region in grayling, suggesting either loss of large duplicated blocks or sequence assembly collapse in grayling. To test the ‘assembly collapse’ hypothesis, we mapped the grayling Illumina paired-end reads that were used for the assembly back to the grayling genome sequence using BWA-MEM (Li 2013) and determined the mapped read depth for each of the grayling genes. Single-copy grayling genes in LORe regions consistently displayed read depths (~100x) twice that of the LORe duplicates in grayling (Figure 2c and Supplementary Figure S4a), indicating assembly collapse rather than loss of large chromosomal regions. Additionally, the SNP density of the scaffolds in these regions computed using FreeBayes (Garrison & Marth 2012) (quality filter of 30) displayed values that were on an average twice that of the background SNP density, albeit with a much wider distribution (Figure 2d and Supplementary Figure S4b).

**Fig. 2.**
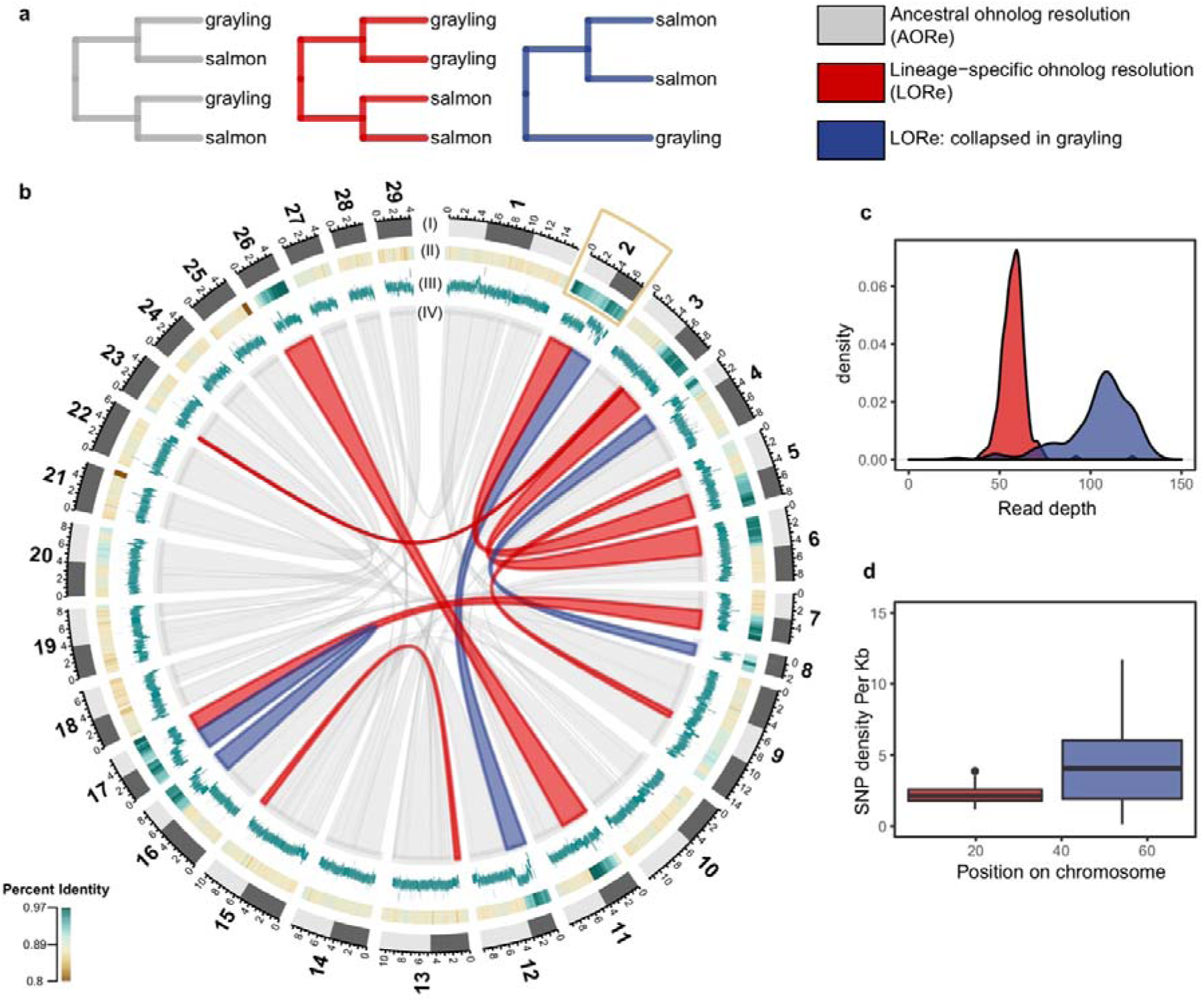
Rediploidization in grayling genome. a) Gene tree topologies corresponding to the different models of ohnolog resolution (ancestral divergence of ohnologs (AORe) and lineage-specific divergence of ohnologs (LORe and LORe-like regions with repeat collapse in grayling). b) Circos plot: Outer track (I) represents the 29 chromosomes of Atlantic salmon with chromosome arms indicated using light and dark grey. (II) Percent identity between duplicated genomic regions in Atlantic salmon with darker green representing higher percent identity (see color scale). (III) Average number of reads mapped to grayling genes in the corresponding regions. (IV) The grey ribbons represent the ancestrally diverged gene duplicate pairs (AORe), while the red ribbons represent the LORe duplicate pairs and the blue ribbons correspond to LORe regions with a collapsed assembly in grayling. The inset plot shows the distribution of average depth of reads mapped to the grayling genes (c) and SNP density per Kb (d) across chromosome 2 (marked with a yellow box in b).

### Ohnolog tissue gene expression regulation

To investigate the regulatory divergence in tissue expression following Ss4R, we exploited tissue expression atlases of Atlantic salmon and grayling in a co-expression analysis. Individual genes from 5,070 ‘expressed’ ohnolog-tetrads (20,280 genes in total) were assigned to eight ‘tissue dominant’ expression clusters (Methods, Supplementary Figure S5). These co-expression clusters were used to identify ohnolog-tetrads conforming to expectations of expression patterns under five evolutionary scenarios (see Table 2, Figure 3): (I) ancestral ohnolog divergence followed by independent purifying selection in both species, (II) conserved tissue regulation in both species, (III and IV) lineage-specific regulatory divergence of one duplicate. In addition, a fifth (V) scenario where regulation among duplicates are shared within species but different between species, is expected to be common in genomic regions with LORe. Further, the ohnolog-tetrads where three, or all four of the duplicates were in different tissue-expression clusters were grouped into a 6th ‘unclassified’ category.

After applying a gene tree topology-based filtering criterion to the 5,070 ohnolog-tetrads (see Methods), 509 conforming to the expectations of LORe and 3,480 conforming to AORe gene tree topologies were retained for further analyses. Of the five evolutionary scenarios, conserved tissue regulation was the most common (~25%), followed by species-specific divergence of a single duplicates (~11% in Atlantic salmon and ~15% in grayling). Category (I) and (V) were the least common categories (Table 2), and as expected, category (V) was significantly enriched in LORe regions (Fisher’s exact test, two-sided, p-value < 0.0005).

**Fig. 3.**
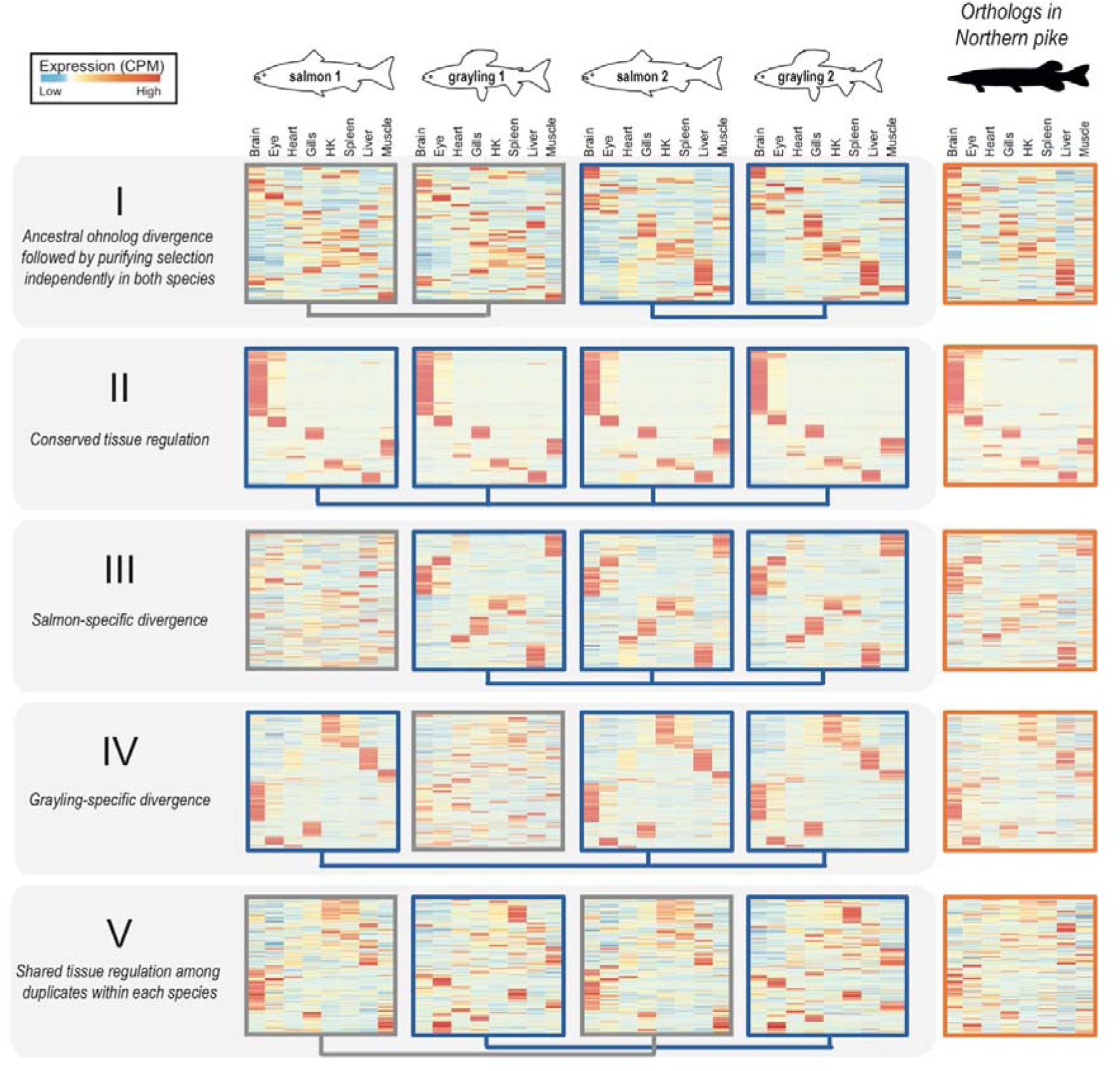
Selection on tissue expression regulation after whole genome duplication. Heatmaps show clustering of tissue expression of the ohnolog-tetrads for each of the five evolutionary scenarios of tissue expression regulation following Ss4R WGD (see Table 2). Within each category, a row across the first four heatmaps in represents one ohnolog-tetrad (four genes: salmon1, grayling1, salmon2 and grayling2), that were ordered based on similarity of expression profiles with the corresponding orthologs in Northern pike (the fifth heatmap depicted on the right). Darker red corresponds to the highest expression level observed for one gene, and the darker blue to the lowest (scaled counts per million, CPM). Connecting blue lines below the heatmaps indicate duplicates belonging to the same tissue clusters (conserved expression pattern). (An extended figure with ohnolog-tetrads subdivided into LORe and AORe in Supplementary Figure S6)

**Table 2:** Classification of tissue expression divergence in the ohnolog-tetrads. The number and percentages of genes in each category calculated based on the total number of topology-filtered ohnolog-tetrads.

| **Evolutionary scenario** | **AORe** | **LORe** |
| --- | --- | --- |
| **(I)** *Ancestral ohnolog divergence followed by purifying selection independently in both species* | 199 (5.7%) | 24 (4.7%) |
| **(II)** *Conserved tissue regulation of all ohnologs.* | 869 (25%) | 131 (25.7%) |
| **(III)** Atlantic salmon-specific divergence of one ohnolog | 375 (10.8%) | 70 (13.8%) |
| **(IV)** Grayling-specific divergence of one ohnolog | 516 (14.8%) | 80 (15.7%) |
| **(V)** *Conserved tissue regulation among ohnologs within species but different between species* | 195 (5.6%) | 51 (10.0%) |
| **Unclassified:** *Ohnolog-tetrads with neutral-like expression evolution* | 1326 (38.1%) | 153 (30.1%) |
| **Total** | 3480 | 509 |

To assess the directionality of the expression divergence relative to the presumed ancestral state, we compared tissue expression of the ohnolog-tetrads to that of the corresponding orthologs in Northern pike (Figure 3). Previous studies have shown that genome-wide tissue-specific expression divergence among WGD ohnologs in teleosts mostly evolve through asymmetric divergence in tissue regulation (Lien et al. 2016; Sandve et al. 2017). The predominant expression pattern thus reflects one ohnolog copy retaining more regulatory similarity with unduplicated orthologs (regulatory neo-functionalization), with very small proportion (<1%) of ohnologs displaying characteristics of regulatory sub-functionalization (Lien et al. 2016). Under a model of sub-functionalization, the sum of expression levels of both ohnologs should correlate better to the assumed ancestral expression regulation than any of the individual ohnologs (Sandve et al. 2017). Therefore, we tested if the divergence patterns leading to maintenance of the two ohnolog copies in the category (I) tetrads are associated with this atypical mode of expression divergence. Both the distribution of ohnolog tissue expression correlations (Supplementary figure S7), and the patterns in the heatmaps (Figure 3), support the regulatory neo-functionalization pattern for all three evolutionary scenarios where we observe ohnolog divergence (category I, III and IV).

As different tissues are involved in different biological functions, we expect that the regulatory evolution is shaped by tissue-specific selective pressures (Gu & Su 2007). To test this, we evaluated the hypothesis that tissues are disproportionately contributing to ohnolog-tetrad divergence by comparing the “tissue dominant” cluster distribution across all tetrads. For all evolutionary scenarios, between 2–5 tissue-expression clusters were significantly over- or underrepresented (Fisher’s tests, two sided, Bonferroni corrected p-value<=0.05), with the conserved category being the most skewed in tissue representation with a strong bias towards brain-specific expression (Supplementary Table S4). The high tissue-specificity (Tau score) of genes in ohnolog-tetrads associated with these genes (Supplementary Figure S8) corroborates the observed brain-specific expression bias.

Further, we tested whether distinct ohnolog-tetrad divergence categories were coupled to patterns of protein-coding and promoter sequence evolution. Specifically, we tested the hypothesis that conserved regulation is associated with conserved protein coding evolution. We estimated the dN/dS ratios for each duplicate pair within each species and compared the distribution of dN/dS statistics in each class with that of the neutral-like (‘Unclassified’) category (Supplementary Figure S9). Low dN/dS (<<1) indicates strong purifying selection pressure. Categories I-V show variability in among-ohnolog dN/dS ratios, with category (I) having significantly higher dN/dS ratio compared to the ‘Unclassified’ category (Wilcoxon rank sum, p=0.005) and categories (II) and (V) having significantly lower dN/dS ratios (Wilcoxon rank sum, p=0.014 and p=0.0017, respectively). The ohnolog pairs showing lineage-specific expression divergence (III and IV) did not have a significantly different dN/dS ratio compared to the neutral-like category (Wilcoxon rank test, p-values=0.36 and 0.26, respectively). Further, we used the Atlantic salmon genome to annotate and compare known transcription factor motifs divergence in the promoters (from 1000 bp upstream to 200 bp downstream of the transcription start site) of ohnologs. Under the assumption that expression divergence is, at least partly, driven by changes in transcription factor binding motifs located in proximal promoters, we tested if ohnolog regulatory divergence in salmon (scenario (I) and (III)) was associated with divergence of promoter motifs. Indeed, the results add validation to the different expression evolution classifications (Supplementary Figure S10), with categories (I) and (III) having significantly less similar promoter motif content compared to ohnolog-tetrads with conserved tissue expression regulation in salmon (II, IV and V) (Wilcoxon test all contrasts between I/III and II/IV/V, p<0.04–0.002).

To evaluate if the ohnologs in different classes were associated with distinct biological functions, we performed GO term enrichment tests. The ohnolog-tetrads of category (II) under strict selective constraints show highly brain-specific expression and are enriched for GO functions related to behaviour and neural functions. In contrast, genes in category (I), which represents ohnologs that underwent divergence in gene regulation following WGD, are associated with functions related to lipid metabolism, development, and immune system (Supplementary file 2).

Highly connected genes in protein-protein interaction networks are often placed under strong constraints to maintain stoichiometry (Freeling & Thomas 2006; Sémon & Wolfe 2007). To test if the strong constraints on the ohnolog-tetrads with conserved tissue expression (II) are associated with having higher protein-protein interactions, we extracted all the zebrafish genes from the genes trees corresponding to the ohnologs in the expression divergence categories and queried them against the STRING database (Szklarczyk et al. 2017) (version 10.5). Only associations with a score of above 7.0, suggesting high confidence associations, were retained. As expected, we found that category (II) genes were indeed enriched for PPI (enrichment p-value, 1.05e-05) in comparison to the genes in the other classes (I, III and IV) with diverged expression (enrichment p-value, 0.785).

### Evolution of gene expression levels following WGD

The co-expression analyses leverage gene expression variation between tissues to classify ohnologs according to regulatory divergence. However, it is important to note that it does not explicitly test for significant changes in gene expression levels. To assess the divergence of ohnolog expression levels, we generated an RNA-Seq dataset from additional liver samples from Atlantic salmon (n=4) and grayling (n=4). We tested for differences in liver expression levels between ohnolog pairs within both species and reported absolute differences in fold change (FC) and the statistical significance (FDR adjusted p-values) for these tests. Of the 2,467 ohnolog-tetrads in categories I-V (Table 2), 54% (1,349) showed significant (FDR<10^−3^) fold change differences (FC>2) in liver expression in at least one species, with 19% (467) in both species, 18% (455) in grayling only, and 17% (427) in salmon only.

We then focussed on the subset of ohnolog-tetrads where at least one ohnolog was assigned to an expression cluster displaying dominant expression in liver. From this subset of 552 ohnolog-tetrads, 80% (442) showed significant (FDR<10^−3^) fold change differences (FC>2) in liver expression in at least one species, 37% (204) both species, 25% (136) grayling only, and 18% (102) salmon only (Supplementary Table S5). As tissue-dominance is the main factor in the analyses of tissue-regulatory evolution, we expected that the different evolutionary scenarios (Figure 3) should be associated with enrichments in certain patterns of expression level divergence, or alternatively, a lack thereof. We indeed found that the ohnolog-tetrads reflecting ancestral divergence followed by purifying selection in both species (scenario I) were significantly enriched in ohnologs being expressed at different levels in both species compared to other scenarios (Figure 4a, Fisher’s test p-value=1.741×10^−4^). Conversely, those ohnolog pairs that show shared tissue regulatory patterns (scenarios II and V) were significantly enriched in ohnologs with no expression level divergence (p-values 0.0117 and 6.79×10^−3^). Finally, ohnologs with tissue regulatory divergence in one species (scenario III and IV in Figure 3) also had the most pronounced enrichment of expression level divergence for that species (Figure 4a, p-values 3.324×10^−7^ and 2.028×10^−6^). Three examples of putative liver-specific expression gains showing high correspondence between tissue regulation and expression level evolution are highlighted in Figure 4c-h.

Further, we assessed if liver-specific expression level differences between ohnologs were associated with changes in transcription factor binding motif presence in promoters. We partitioned the ohnologs into three categories; differentially expressed in both species (likely diverged in expression in an ancestor of all salmonids), species-specific expression level divergence (we only used salmon-specific cases as we had no promoter motif data for grayling), and no significant difference in expression level. The lowest promoter motif similarity was found among ohnologs where both species showed strong expression level divergence, followed by the salmon-specific and then no expression divergence (Figure 4b).

**Fig. 4.**
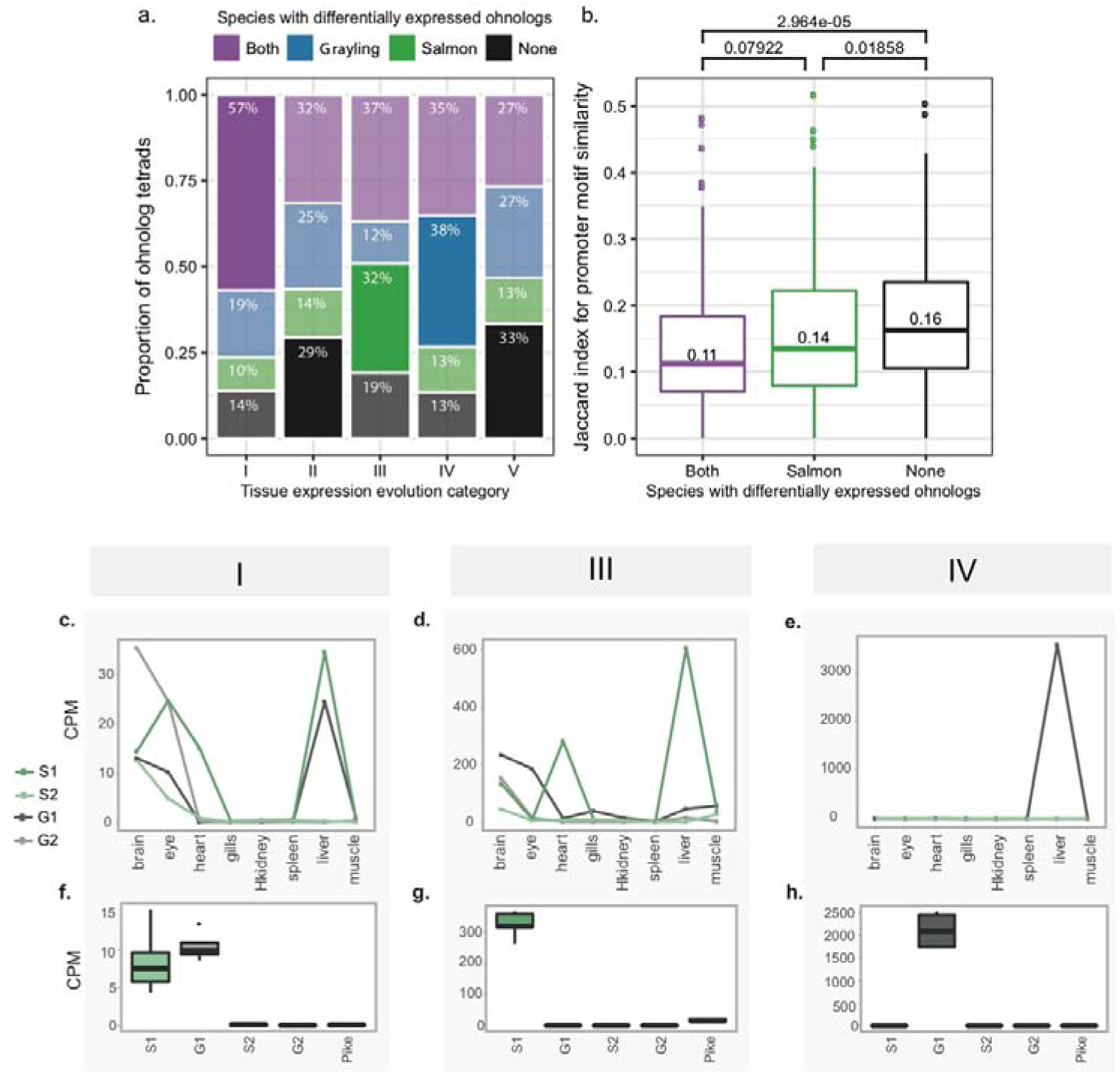
Expression level evolution of the ohnolog-tetrads. (a-b) Ohnolog-tetrads were tested using liver expression data for cases of highly significant differential expression (FDR adjusted p-value <10^−3^, absolute fold change >2) between ohnologs of both species (purple), grayling ohnologs only (blue), Atlantic salmon ohnologs only (green), and no onhologs of either species (black). Ohnolog tetrads shown had at least one ohnolog assigned to a tissue expression cluster displaying dominant expression in liver. a) The number and proportions of cases are given per tissue expression evolution category (see Table 2). The differential expression outcome expected to be the highest for each category is highlighted opaque, while the rest are transparent. b) Jaccard index score distributions for Atlantic salmon ohnolog promoter motif similarity, separated by differential expression outcome. P-values from pairwise comparisons testing for lower Jaccard index using the Wilcoxon test are indicated - as well as the median scores. (c-h) Expression levels, in terms of CPM (counts per million), from the tissue atlas data (c-e) and the corresponding data from the liver expression data are plotted (boxplots in f-h) for a selected example with a liver-specific gain of expression in each of the categories (I), (III) and (IV). The examples indicated in (c-h) include ohnologs of ephrin type-B receptor 2-like (category I), contactin-1a-like gene (category III) and a E3 ubiquitin-protein ligase-like gene (category IV). The ohnologs in Atlantic salmon and grayling are represented as S1, S2 and G1 and G2 respectively.

### Loss of purifying selection on chloride ion transporter regulation in non-anadromous grayling

The most apparent difference in biology between grayling and Atlantic salmon is the ability of Atlantic salmon to migrate between freshwater and saltwater (anadromy), a trait that grayling has not evolved. Saltwater acclimation involves changes in switching from ion absorption to ion secretion to maintain osmotic homeostasis. To assess whether key genes associated with the ability to adapt to seawater are under divergent selection for expression regulation in Atlantic salmon and grayling, we probed into categories (III) and (IV) for overrepresented GO terms related to ion-homeostasis (i.e. potassium, sodium or chloride regulation/transport). Interestingly, in the group exhibiting grayling-specific regulatory divergence (category IV), we found that ‘regulation of chloride transport’ was overrepresented. One of the genes associated with this GO term was the classical anadromy-associated, salinity-induced, cystic fibrosis transmembrane conductance regulator (CFTR). The CFTR gene transports chloride ions over cell membranes in the gill and is involved in saltwater adaptations in Atlantic salmon (Nilsen et al. 2007). Looking into the tissue expression profiles of this tetrad (Figure 5a) it was evident that the divergence of tissue regulation in grayling was associated with a loss of gill tissue expression specificity compared to Atlantic salmon. To determine if the grayling CFTR duplicate with diverged expression also had signatures of coding sequence divergence, we computed branch-specific dN/dS. Notably, the grayling CFTR displaying diverged expression regulation also displays a two-fold increase in dN/dS compared to its Ss4R duplicate with conserved expression regulation, reflecting relaxation of purifying selection pressure on one ohnolog in the non-anadromous grayling (Figure 5b). These results may suggest that there could be a fitness advantage of maintaining two copies of gill expressed CFTR for anadromous species, but not for pure freshwater adapted species, such as grayling.

**Fig. 5.**
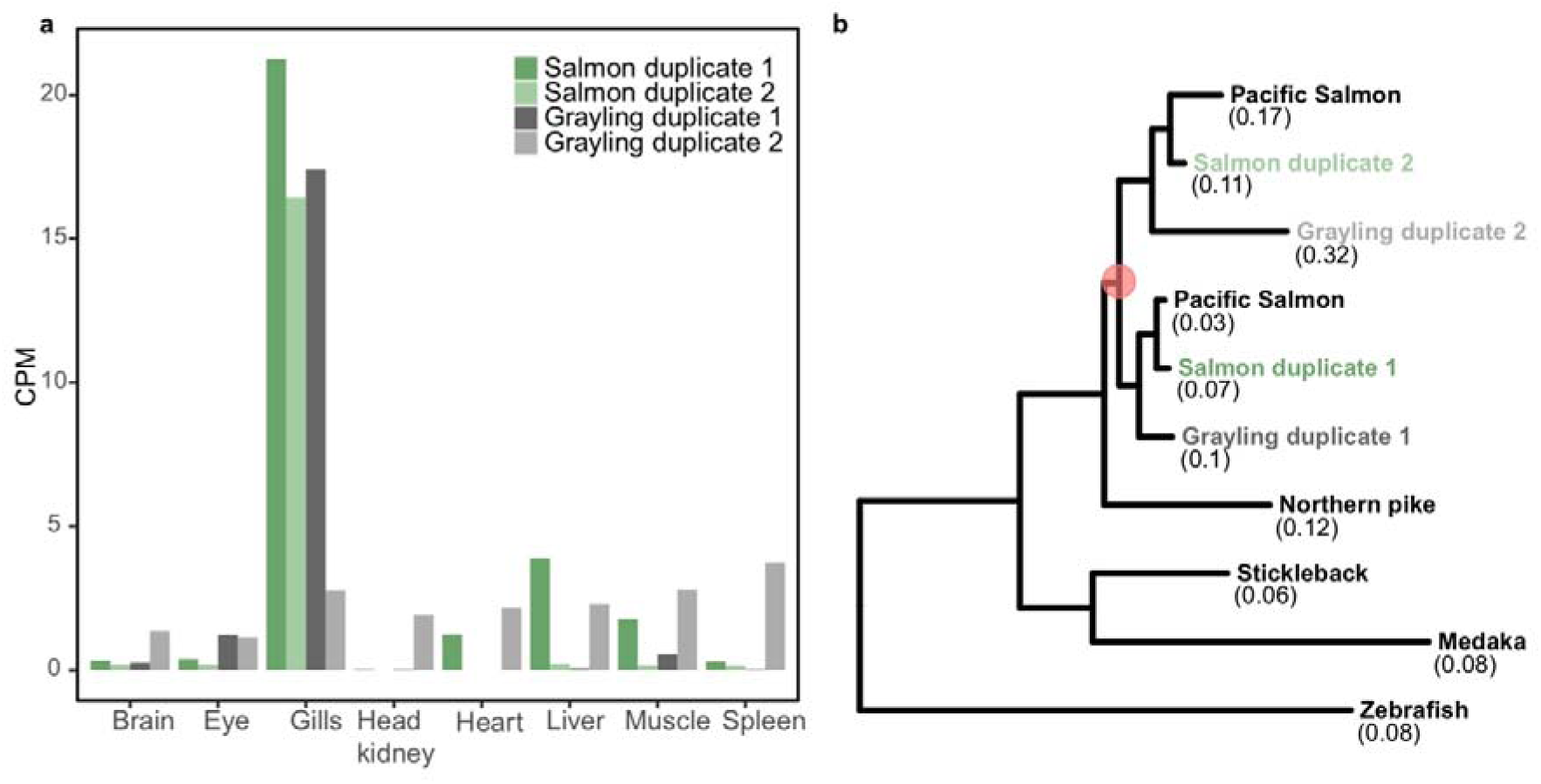
Divergent selection on cystic fibrosis transmembrane conductance regulator. a) Expression values, in terms of counts per million (CPM), of the cystic fibrosis transmembrane conductance regulator (CFTR) ohnologs in Atlantic salmon and grayling across eight tissues. b) CFTR gene tree. The orange circle represents the Ss4R duplication. Branch-specific dN/dS of the tip nodes are given in parentheses.

## Discussion

A major limitation in previous studies of evolution of gene regulation following WGD in vertebrates has been the inability to distinguish between neutral and adaptive divergence (Hermansen et al. 2016). Here, we leverage gene expression data from two salmonid species and a close outgroup species (Northern pike) in a comparative approach to identify shared expression evolution patterns following WGD in lineages evolving independently for ~50 million years. This allows us to identify evolutionarily long-term conservation of novel expression “phenotypes” arising after WGD - the hallmark of novel adaptive functions. Although regulatory divergence of Ss4R ohnologs is widespread (Lien et al. 2016; Gillard et al. 2018), we show that ohnolog regulatory tissue divergence shared among species separated by ~50 million years of evolution is comparably rare (Table 2). Nevertheless, these ohnologs represent intriguing candidates for salmonid-specific adaptive evolution of novel gene expression regulation following WGD. Salmonids are suggested to have evolved from a pike-like ancestor; a relatively stationary ambush predator (Craig 2008). Under this assumption, early salmonid evolution must have involved adaptation to new pelagic and/or riverine habitats. Adaptations to new environments and evolution of different life history strategies are known to be associated with strong selective pressure on immune-related genes (e.g: (Haase et al. 2013; Solbakken et al. 2017)). In line with this, we see an overrepresentation of immune-related genes among ohnologs that diverged in the common ancestor of salmon and grayling but have been under purifying selection in both species after speciation (category I, Table 2). Furthermore, pike is generally piscivorous throughout their lifespan, while salmonids depend more on aquatic and terrestrial invertebrate prey with significantly lower input of lipids, especially in early life (Carmona-Antoñanzas et al. 2013). Interestingly, duplicates with shared ancestral divergence (category I), which are candidates for adaptive divergence in regulation, are enriched for liver-expressed genes involved in lipid-homeostasis, metabolism and energy storage (glycogen) related functions (GO test results in Supplementary file 2). Taken together, our results suggest a role of Ss4R ohnologs in adaptive evolution of novel gene expression regulation, possibly related to new pathogenic pressures in a new type of habitat, and optimization of lipid-homeostasis and glycogen metabolism-related functions in response to evolution of a more active pelagic/riverine life with limited lipid resources.

Purifying selection to maintain ancestral tissue regulation of ohnologs in both salmonid species was the most commonly observed fate of ohnolog expression evolution (category II, Table 2 and Figure 3). These ohnologs were predominantly brain-specific and enriched for predicted protein-protein interactions. Several other studies in vertebrates have found similar results, with strong purifying selection on sequence and expression evolution in brain-dominant genes, as well as high retention probability of following large-scale genome duplication (Zheng-Bradley et al. 2010; Chan et al. 2009; Khaitovich et al. 2005; Roux et al. 2017; Guschanski et al. 2017). As neuron-function related genes are involved in complex networks of signalling cascades and higher-order protein-protein interactions, this pattern is believed to be driven either by direct selection for maintaining novel gene copies due to dosage balance (relative or absolute), or indirectly, through selection against toxic effects of misfolding or mis-interacting proteins (Roux et al. 2017).

Recent analyses of salmonid genomes have revealed ~25% LORe between Atlantic salmon and grayling (Robertson et al. 2017; Lien et al. 2016). Here we find a set of LORe regions, corresponding to whole chromosome arms in Atlantic salmon, detected as single copy genes in grayling as a result of collapse during the assembly process. This strongly suggests that these sequences are in fact present as near-identical duplicated regions in the grayling genome. The larger chromosome arm-sized regions virtually indistinguishable at the sequence level (~10% in total, i.e. blue ribbons in Figure 2b) are likely still recombining or have only ceased to do so in the recent evolutionary past. Large-scale chromosomal rearrangements often follow genome duplication to block or hinder recombination among duplicated regions (Comai 2005; Lien et al. 2016). The difference we observe in the rediploidization history between Atlantic salmon and grayling is thus likely linked to the distinctly different chromosome evolution in these species (Supplementary Figure S1) (Qumsiyeh 1994).

The genomic footprints of LORe also extend to ohnolog regulatory divergence. LORe regions showed a strong enrichment of species-specific tissue-specific expression pattern (category V, in Table 2 and Supplementary Table S5), as expected under lineage-specific rediploidization and subsequent regulatory divergence. However, we also find a small proportion (~5%) of category (V) genes in AORe regions of the genome. This observation is more difficult to explain, but it is likely a real biological observation as the coding- and promoter sequence evolution analyses (Supplementary figure S9 and S10) support the validity of the category V ohnologs in AORe regions. Possible explanations for this observation could be a result of non-homologous gene conversion (e.g. (Hastings 2010)) or alternatively, rare local lineage-specific non-homologous recombination events outside the LORe regions.

Finally, one fundamental difference between European grayling and Atlantic salmon is that Atlantic salmon has the ability to migrate between fresh- and seawater (anadromy). In line with this we found that the ohnologs of CFTR, a key gene regulating chloride ion export in gills (Marshall & Singer 2002)involved in saltwater adaptation in Atlantic salmon (Nilsen et al. 2007), seems to be under divergent selection pressure in the two species. One ohnolog in grayling had lost tissue specificity and gill expression dominance and evolved under relaxed purifying selection at the coding sequence level (Figure 5a and b). Atlantic salmon on the other hand have retained both copies as “gill specific”. We thus propose that maintaining two functional CFTR genes could be an adaptive trait in anadromous salmonids in that it improves their ability to remove excess chloride ions and maintain ion homeostasis in the sea. Conversely, in non-anadromous species, there is no selective pressure to maintain both CFTR copies, and this has resulted in the return to a single functional CFTR ohnolog copy in grayling.

## Conclusions

We present a draft genome assembly of European grayling and use it for comparative studies with the reference genome assembly of Atlantic salmon. Our comparative genome and transcriptome analysis between Atlantic salmon and grayling provides novel insights into evolutionary fates of ohnologs subsequent to WGD and associations between signatures of selection pressures on gene duplicate regulation and the evolution of salmonid traits, including anadromy. Hence, the genome resource of grayling opens up new exciting avenues for utilizing salmonids as a model system to understand the evolutionary consequences of WGD in vertebrates.

## Methods

### Sampling and sequencing

A male grayling specimen was sampled outside of its spawning season (October 2012) from the River Glomma at Evenstad, Norway. The fish was humanely sacrificed and various tissue samples were immediately extracted and conserved for later DNA and RNA analysis. Fin clips were stored on 96% ethanol for DNA sequencing. Tissues from muscle, gonad, liver, head kidney, spleen, brain, eye, gill and heart were stored in RNAlater for RNA extraction. The DNA was extracted from fin clips using a standard high salt DNA extraction protocol. A paired-end library with an insert size ~180 (150 bp read length) and mate pair libraries of insert size ~3kb and 6 kb (100bp read length) were sequenced using the Illumina HiSeq2000 platform (Table S1). Total RNA was extracted from the different tissue samples using the RNeasy mini kit (Qiagen) following the manufacturer’s instructions. The library construction and sequencing were carried out using Illumina TruSeq RNA Preparation kit on Illumina HiSeq2000 (Table S2). All the library preparation and sequencing were performed at the McGill University and Génome Québec Innovation Centre.

### Genome assembly and validation

The sequences were checked for their quality and adapter trimming was performed using cutadapt (version 1.0) (Martin 2011). A *de novo* assembly was generated with Allpaths-LG (release R48777) (Gnerre et al. 2011) using the 180bp paired-end library and the mate pair (3kb and 6kb) libraries. Assembly polishing was carried out using pilon (version 1.9) (Walker et al. 2014). The high copy number of mitochondrial DNA often leads to high read coverage and thus misassembly. The mitochondrial genome sequence in the assembly was thus reassembled by extracting the reads that mapped to the grayling *(Thymallus thymallus)* mtDNA sequence (GenBank ID: NC_012928), followed by a variant calling step using Genome Analysis Toolkit (GATK) (version 3.4–46) (Van der Auwera et al. 2013). The consensus mtDNA sequence thus obtained was added back to the assembly.

To identify and correct possibly erroneous grayling scaffolds, we aligned the scaffolds against a repeat masked version of the Atlantic salmon genome (Lien et al. 2016) using megablast (E-value threshold 1e-250). Stringent filtering of the aligned scaffolds (representing 1.3 Gbp of the 1.4 Gbp assembly) identified 13 likely chimeric scaffolds mapping to two or more salmon chromosomes (Supplementary File 1), which were then selectively ‘broken’ between, apparently, incorrectly linked contigs.

### Transcriptome assembly

The RNA-Seq data from all the tissue samples were quality checked using FastQC (version 0.9.2). The sequences were assembled using the following two methods. Firstly, a *de-novo* assembly was performed using the Trinity (version 2.0.6) (Grabherr et al. 2011) pipeline with default parameters coupled with *in silico* normalization. This resulted in 730,471 assembled transcript sequences with a mean length of 713 bases. RSEM protocol-based abundance estimation within the Trinity package was performed where the RNA-Seq reads were first aligned back to the assembled transcripts using Bowtie2 (Faust & Hall 2012), followed by calculation of various estimates including normalized expression values such as FPKM (Fragments Per Kilobase Million). A script provided with Trinity was then used to filter transcripts based on FPKM, retaining only those transcripts with a FPKM of at least one. Secondly, reference-guided RNA assembly was performed by aligning the RNA reads to the genome assembly using STAR (version 2.4.1b) (Dobin et al. 2013). Cufflinks (version 2.1.1) (Dobin et al. 2013; Trapnell et al. 2010) and TransDecoder (Haas et al. 2013) were used for transcript prediction and ORF (open reading frame) prediction, respectively. The resulting transcripts were filtered and retained based on homology against zebrafish and stickleback proteins, using BlastP and PFAM (1e-05). The *de-novo* method resulted in 134,368 transcripts and the reference-based approach followed by filtering resulting in 55,346 transcripts.

### Genome Annotation

A *de novo* repeat library was constructed using RepeatModeler with default parameters. Any sequence in the *de-novo* library matching a known gene was removed using Blastx against the UniProt database. CENSOR and TEclass were used for classification of sequences that were not classified by RepeatModeler. Gene models were predicted using an automatic annotation pipeline involving MAKER (version2.31.8), in a two-pass iterative approach (as described in https://github.com/sujaikumar/assemblage/blob/master/README-annotation.md). Firstly, *ab initio* gene predictions were generated using GeneMark ES (version 2.3e) (Lomsadze 2005) and SNAP (version 20131129) (Korf 2004) trained on core eukaryotic gene dataset (CEGMA). The first round of MAKER was then run using the thus generated *ab initio* models, with the UniProt database as the protein evidence, the *de novo* identified repeat library and EST evidences from the transcriptomes assembled using *de novo* and the reference guided approaches, along with the transcript sequences from the recent Atlantic salmon annotation (Lien et al. 2016). The second pass involved additional data from training AUGUSTUS (Stanke et al. 2008) and SNAP models on the generated MAKER predictions.

Putative functions were added to the gene models using BlastP against the UniProt database (e-value 1e-5) and the domain annotations were added using InterProScan (version 5.4–47) (Quevillon et al. 2005). Using the MAKER standard filtering approach, the resulting set of genes were first filtered using the threshold of AED (Annotation Edit Distance), retaining gene models with AED score less than 1 and PFAM domain annotation. AED is a quality score given by MAKER that ranges from 0 to 1 and indicates the concordance between predicted gene model and the evidence provided, where an AED of 0 indicates that the gene models completely conforms to the evidence. Further, for the genes with AED score of 1 and no domain annotations, a more conservative Blast search was performed against UniProt proteins and Atlantic salmon proteins with an e-value cut-off of 1e-20. The genes with hits to either of these databases were also retained. The completeness of the annotations was again assessed using CEGMA (Parra et al. 2007) and BUSCO (Simão et al. 2015).

### Analysis of orthologous groups

We used orthofinder (version 0.2.8, e-value threshold at 1e-05) (Emms & Kelly 2015) to identified orthologous gene groups (i.e orthogroup). As input to orthofinder, we used the MAKER-derived *T. thymallus* gene models as well as protein sequences from three additional salmonid species (Atlantic salmon, Rainbow trout and coho salmon), four non-salmonid teleost species *(Esox lucius, Danio rerio, Gasterosteus aculeatus, Oryzias latipes)*, and two mammalian outgroups *(Homo sapiens, Mus musculus)*. Rainbow trout protein annotations were taken from https://www.genoscope.cns.fr/trout/. Atlantic salmon, *Esox lucius* data were downloaded from NCBI ftp server (ftp://ftp.ncbi.nih.gov/genomes/, release 100). The transcriptome data for Coho salmon was obtained from NCBI (GDQG00000000.1) and translated using TransDecoder. All other annotations were downloaded from ENSEMBL. Each set of orthogroup proteins were then aligned using MAFFT(v7.130) (Katoh et al. 2002) using default settings and the resulting alignments were then used to infer maximum likelihood gene trees using FastTree (v2.1.8) (Price et al. 2010) (Figure 1 a and b). As we were only interested in gene trees containing information on Ss4R duplicates, complex orthogroup gene trees (i.e. containing 2R or 3R duplicates of salmonid genes) were subdivided into the smallest possible subtrees. To this end, we developed an algorithm to extract all clans (defined as unrooted monophyletic clade) from each unrooted tree (Wilkinson et al. 2007) with two monophyletic salmonid tips as well as non-salmonid outgroups resulting in a final set of 20,342 gene trees. In total, 31,291 grayling genes were assigned to a clan (Figure 1 and Supplementary Figure S2). We then identified homoelogy in the Atlantic salmon genome by integrating all-vs-all protein BLAST alignments with a priori information of Ss4R synteny as described in Lien et al. 2016 (Lien et al. 2016). Using the homeology information, we inferred a set of high confidence ohnologs originating from Ss4R. The clans were grouped based on the gene tree topology into duplicates representing LORe and those with ancestrally diverged duplicates (AORe). The LORe regions were further categorized into two (duplicated or collapsed) based on the number of corresponding *T.thymallus* orthologs. This data was plotted on Atlantic salmon chromosomes using circos plot generated using OmicCircos (https://bioconductor.org/packages/release/bioc/html/OmicCircos.html). The LORe and AORe ohnologs with two copies in each species are hereafter referred to as ohnolog-tetrads.

### Expression divergence and conservation

The grayling RNA-Seq reads from each of the eight tissues (liver, muscle, spleen, heart, head kidney, eye, brain, gills) were mapped to the genome assembly using STAR (version 2.4.1b). The reads uniquely mapping to the gene features were quantified using htseq-count (Anders et al. 2015). The CPM value (counts per million), here used as a proxy for expression, was then calculated using edgeR (Robinson et al. 2010). Similar CPM datasets were obtained from Atlantic salmon RNA-seq data reported in Lien et al. (Lien et al. 2016).

Filtering of ortholog groups (i.e. clans) was performed prior to analyses of expression evolution of Ss4R ohnologs: 1) we only considered Ss4R duplicates that were retained in both Atlantic salmon and grayling, 2) the Ss4R duplicates were classified into AORe or LORe, based on topologies of the ortholog group gene trees, only gene pairs with non-zero CPM value were considered. This filtering resulted in a set of 5,070 duplicate pairs from both Atlantic salmon and grayling (ohnolog-tetrads). The gene expression values from the gene duplicates in the ohnolog-tetrads were clustered using hclust function in R, using Pearson correlation into eight tissue dominated clusters. The expression pattern in the eight clusters of the genes in ohnolog-tetrads was used to further classify them into one of the ohnolog expression evolution categories (see Table 2). The ohnolog-tetrads were further filtered based on expected topologies under LORe and AORe scenarios. Heatmaps of expression counts were plotted using pheatmap package in R (https://CRAN.R-project.org/package=pheatmap). To quantify the breadth of expression (i.e., the number of tissues a gene is expressed in), we calculated the tissue specificity index Tau (Yanai et al. 2005) for all the genes in ohnolog-tetrads, where a τ value approaching 1 indicates higher tissue specificity while 0 indicates ubiquitous expression.

### Expression comparison in liver

Utilizing independent liver tissue samples, we compared differential expression in in liver tissue gene expression among ohnologs of grayling and Atlantic salmon with their ohnolog-tetrad tissue expression evolution categories. The liver samples from four grayling individuals were sampled in the river Gudbrandsdalslågen. The samples were from two males (370, 375 mm) and two females (330, 360 mm). The fish was euthanized and dissected immediately after capture and the liver was stored in RNAlater. Total RNA was extracted and 100bp single-end read libraries were generated for two individuals and sequenced using using the Illumina HiSeq4000 platform. For the other two individuals, 150bp paired-end read libraries were generated and sequenced using the Illumina HiSeq2500 platform. RNA-seq data for an additional four Atlantic salmon liver tissue samples was obtained from a feeding experiment (Gillard et al. 2018). Pre-smolt salmon were raised on fish oil-based diets under freshwater conditions.

The RNA-Seq read data was quality processed using CutAdapt (Martin 2011) before alignment to grayling or Atlantic salmon (ICSASG_v2, (Lien et al. 2016)) genomes respectively using STAR (Dobin et al. 2013). RSEM (Li & Dewey 2011) expected counts were generated for gene features. EdgeR (Robinson et al. 2010) was used to generate normalized library sizes of samples (TMM normalization), followed by a differential expression analysis using the exact test method between the gene expression of both the grayling and Atlantic salmon ohnologs in each ohnolog-tetrad. The fold change (log2 scaled) and significance of differential expression (false discovery rate corrected p-values) were produced for grayling and Atlantic salmon duplicates, as well as relative counts in the form of CPM.

### Sequence evolution

To estimate coding sequence evolution rates, we converted amino acid alignments to codon alignments using pal2nal (Suyama et al. 2006). The *seqinr* R package (http://seqinr.r-forge.r-project.org/) was used to calculate pairwise dN and dS values for all sequences in each alignment using the *“kaks”* function. For in-depth analyses of branch specific sequence evolution of the CFTR genes, we used the codeml in PAML (version 4.7a) (Yang 1997). To assess if sequences in the CFTR gene tree evolved under similar selection pressure we contrasted a fixed dN/dS ratio (1-ratio) model and a free-ratio model of codon evolution. A likelihood ratio test was conducted to assess whether a free ratio model was a significantly better fit to the data. Branch specific dN/dS values were extracted from the ML results for the free ratios model.

The two Pacific salmon genes in the CFTR tree (Figure 5) correspond to a gene from Rainbow trout and another from Coho salmon. A blat search of CFTR gene against the Rainbow trout assembly (https://www.genoscope.cns.fr/trout/) resulted in hits on three different scaffolds, with one complete hit and two other partial hits on unplaced scaffolds. Additionally, Coho salmon data is based on a set of genes inferred from transcriptome data. Therefore, the presence of a single copy in the tree for the two species is likely an assembly artefact.

### Genome-wide identification of transcription factors binding sites

A total of 13544 metazoan transcription factor protein sequences together with their binding site represented as position specific scoring Matrices (PSSMs referred to as motifs) were collected from transcription factor binding profile databases such as CISBP, JASPAR, 3D-footprint, UniPROBE, HumanTF, HOCOMOCO, HumanTF2 and TRANSFAC^®^.

DNA sequences from upstream promoter regions of Atlantic salmon (−1000bp/+200bp from TSS) were extracted. A first order Markov model was created from the entire set of upstream promoter regions using the fasta-get-markov program in the MEME Suite (Bailey et al. 2009). This background model was used to convert frequency matrices into log-odds score matrices. We performed a genome-wide transcription factors binding sites prediction in the Atlantic salmon genome using the PSSM collection and the Finding Individual Motif Occurrences (FIMO) (Grant et al. 2011) tool in the MEME Suite (p-value = 0.0001 and FDR = 0.2).

Motif similarity between Atlantic salmon ohnolog promoters was scored using the *Jaccard coefficient*. The promoter *Jaccard coefficient* is defined as;

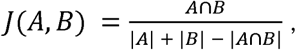
where A and B represents the type of motifs that were present in promoters of the A and B ohnolog copies. If A and B are empty, we set *J(A,B)* = 0 where 0 ≤ *J(A,B)* ≤ 1.

### Gene Ontology (GO) analysis

The gene ontology term (GO) enrichment analysis was performed using the *elim* algorithm implemented in the *topGO* R package (http://www.bioconductor.org/packages/2.12/bioc/html/topGO.html), with a significance threshold of 0.05 using all Ss4R duplicates as the background. GO terms were assigned to salmon genes using Blast2GO (Conesa et al. 2005).

## Availability of data

The Illumina reads have been deposited at ENA under the project accession: PRJEB21333. The genome assembly and annotation data are available at https://doi.org/10.6084/m9.figshare.c.3808162. Atlantic salmon liver expression data is available at ENA or NCBI under sample accessions: SAMEA104483623, SAMEA104483624, SAMEA104483627, and SAMEA104483628.

## Author contributions

KSJ, LAV, SJ and SL conceived and planned the project and generation of the data. SRS and SV performed all the analyses with help from AJN and OKT. Differential expression analysis on the liver dataset was performed by GBG and TRH, and the promoter motif analysis was prepared by TDM. SRS, SV and AJN drafted the manuscript. All authors read and commented on the manuscript.

## Acknowledgements

This work was supported by funding from University of Oslo to the SAK project “Building a marine genome hub” and the Strategic Research Initiative, Center for Computational Inference in Evolutionary Life Science (CELS) to KSJ. SRS was funded partly by the Norwegian Research Council project <u>NFR 244164</u>. We thank Kim M. Bærum for sampling of grayling and Marianne H. S. Hansen for excellent technical assistance. Sample preparation, library construction and sequencing were carried out at the Norwegian Sequencing Centre (NSC), Norway and McGill University and Génome Québec Innovation Centre, Canada. The computational work was performed on the Abel Supercomputing Cluster (Norwegian Metacenter for High-Performance Computing (NOTUR) and the University of Oslo), operated by the Research Computing Services group at USIT, the University of Oslo IT-department. We thank Geir O. Storvik for helpful discussions. We greatly appreciate Daniel J. Macqueen, Marine S. Brieuc and Monica H. Solbakken for critical reading of the manuscript.

